# NMDAR hypofunction increases top-down influence on sensory processing

**DOI:** 10.1101/104588

**Authors:** Adam Ranson, Eluned Broom, Anna Powell, Jeremy Hall

**Affiliations:** Neurosciences & Mental Health Research Institute, Cardiff University, Cardiff CF24 4HQ; School of Biosciences, Cardiff University, Cardiff CF10 3AX; School of Psychology, Cardiff University, Cardiff, CF10 3AT

## Abstract

Converging evidence implicates NMDAR disruption in the pathogenesis of schizophrenia, a condition in which perceptual disturbances are prominent. To explore how NMDAR hypofunction causes perceptual symptoms we investigated activity in cortical sensory circuits in awake behaving mice during pharmacologically induced NMDAR hypofunction. We observed a reduction in sensory-driven activity in V1 while input from the anterior cingulate cortex simultaneously increased, suggesting NMDAR hypofunction may lead to altered perception by modifying the balance of top-down and bottom-up processing.

## Main text

Hypofunction of the N-methyl-D-aspartate receptor (NMDAR) is implicated in the pathogenesis of schizophrenia. Evidence for this association has come both from large scale genetic studies ^1–4^, as well as a large body of experimental work. In humans administration of sub anaesthetic doses of NMDAR antagonists such as ketamine and phencyclidine produce many of the symptoms of schizophrenia including disorganised thought and perceptual distortions^5–7^, and exacerbate symptoms in drug-free schizophrenic patients^8^. Interestingly, global disruption of NMDAR function appears to have differential effects in different cortical regions, with both human imaging and animal studies providing evidence for elevated activity in prefrontal brain regions while other areas such as sensory cortex show unchanged or decreased activation^5,9–13^. This regionally specific disruption of neural activity is relevant in the context of theories of the causes of hallucinations and other positive symptoms of schizophrenia which propose disturbances in transmission of predictive signals from higher to lower levels of the cortical hierarchy (termed ‘top-down’ signals) as a potential cause of perceptual disturbances and in turn delusions^14,15^. One attractive hypothesis is that the perceptual disturbances observed in schizophrenia might be due to an altered balance of influence on of internally generated top-down signals verses bottom-up (sensory) signals on sensory cortex.

In order to directly investigate the effects of global NMDAR antagonism on the integration of bottom-up and top-down signals during sensory processing we used *in vivo* calcium imaging in awake mice (Fig 1a). During ongoing visual stimulation, we measured activity of excitatory and inhibitory neurons in primary visual cortex (V1), and contrasted this with the activity of top-down axons originating from the ACC and measured at their termination site in layer 1 of V1. These axons provide long range top-down feedback to layer 1 of V1, and are thought to mediate attentional and predictive signals^16,17^. Following baseline recordings, top-down ACC axons and V1 activity were then re-measured after either systemic injection of the NMDAR antagonist MK801 or saline control. In order to record V1 activity, neurons were labelled with the genetically encoded calcium indicator GCaMP6S, targeted using intrinsic signal functional imaging (Fig 1b). In addition PV+ interneurons were co-labelled with tdTomato using a PV-cre mouse line (Fig 1c). Activity of top-down projections to V1 from ACC were recorded by transfecting ACC neurons with GCaMP6S and recording their axons at their termination site in layer 1 of V1 (Fig 1d). Recordings were made in awake head-immobilised mice, free to walk on a cylindrical treadmill (Fig 1a; see methods). After a baseline recording and visual stimulation period, animals were administered with either a sub-anaesthetic dose of the NMDAR antagonist MK801 (0.1mg/kg s.c., the previously established optimal dose to elicit increased PFC activity^5,11^) or the same volume of saline and recording was then continued for 45 minutes under identical stimulation conditions. Strikingly, in contrast to reports from other cortical regions (PFC^5,11^; retrosplenial cortex^12^), MK801 treatment reduced V1 population activity by approximately 40% while no reduction in activity was observed after saline injection (Baseline normalised activity: Saline = 0.99 ± 0.096, n = 7 mice, P = 0.23, MK801 = 0.57 ± 0.066, n = 9 mice, P = 0.0002; Fig 1e,f**, Supplementary Video. 1**). In contrast during the same period we observed and approximate 2.5 fold increase in activity of top-down input from ACC while after saline administration no change was observed (Baseline normalised activity: Saline = 1.32 ± 0.275, n = 6 mice, P = 0.3, MK801 = 2.57 ± 0.313, n = 7 mice, P = 0.002; Fig 1g,h**, Supplementary Video. 1**). Consistent with previous freely moving behavioural studies we reliably observed increased locomotor behaviour after MK801 administration (Supplementary Fig. 1a). Locomotion is associated with increased activity in a number of brain regions including frontal cortex and sensory cortex and changes in locomotor behaviour could potentially influence neuronal activity in V1 or ACC. While we observed differential activity changes in V1 (decreased activity) and ACC (increased activity) on MK801 treatment we nevertheless sought to examine the extent to which altered neural activity in general could be accounted for by a change in locomotor behavioural state. To address this question we limited our analysis of V1 and ACC activity to pre and post injection periods of matched locomotor activity (moving periods) and found a similar pattern whereby NMDAR blockade resulted in decreased activity in V1, and increased activity of top down input from ACC (Supplementary Fig. 1b,c) suggesting differences in locomotor behaviour was not in itself the cause of changes observed in neural activity.

**Figure 1:**
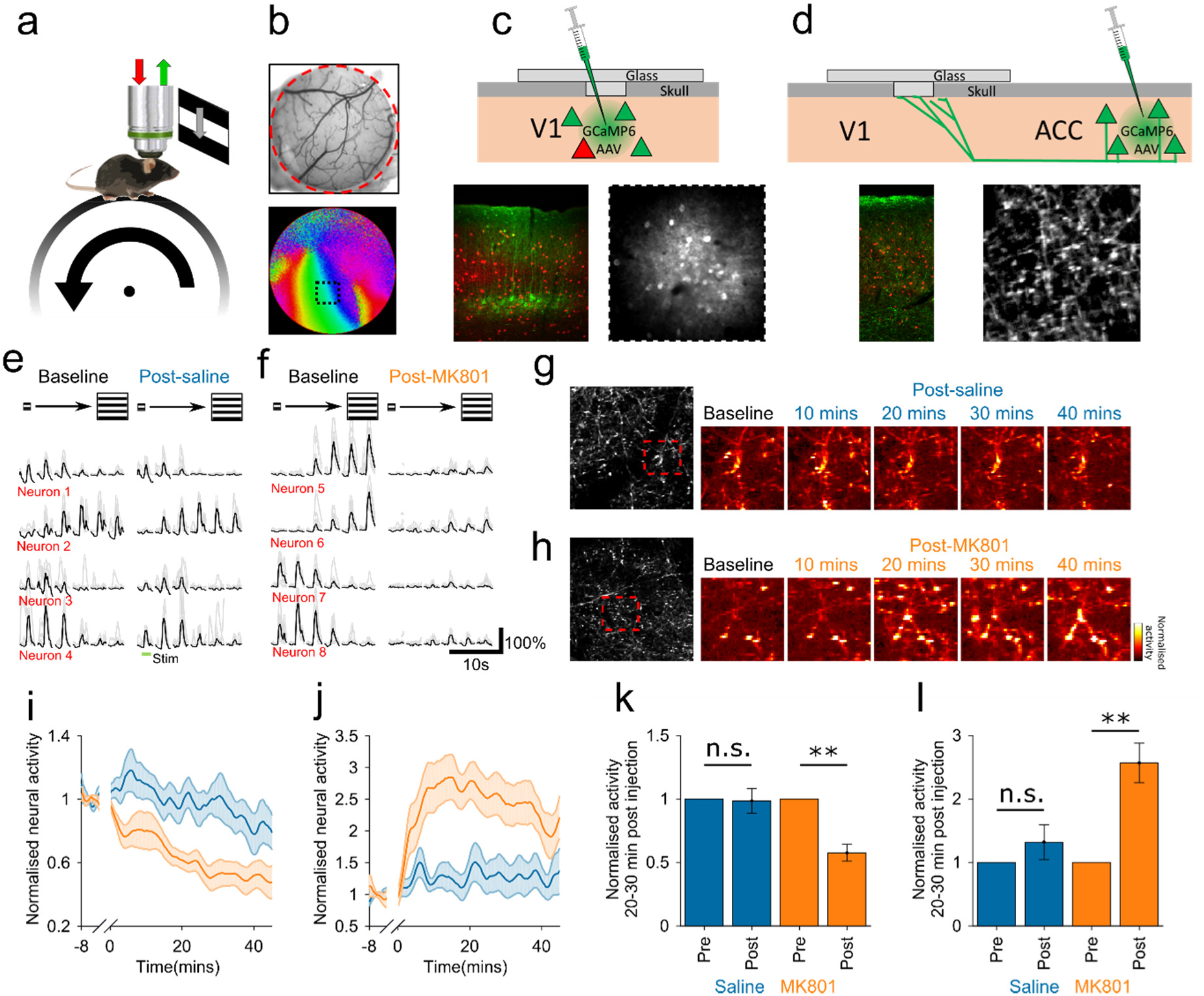
Opposite modulation of sensory verses top down drive to sensory cortex during NMDAR antagonism. (**a**) Schematic of visual stimulation and recording setup. (**b**) Cranial window and intrinsic signal functional map of V1 with representative field of view placement (black square). (**c-d**) Schematic of V1 soma and ACC axon recording configuration, and *ex vivo* and *in vivo* images of recorded tissue. (**e-f**) Examples traces of averaged stimulus evoked responses of single V1 putative excitatory neurons pre and post injection in saline (**e**) and MK801 (**f**) conditions. (**g-h**) Example activity maps of V1eACC axons at baseline and after administration of saline (**g**) or MK801 (**h**). (**i-j**) Normalised V1 somatic population activity (**i**) and V1)ACC axon activity (**j**) after saline (blue) or NMDAR antagonist injection (orange, MK801) during visual stimulation. (**k-l**) Average activity of V1 population (**k**) or ACC axons (**l**) at 20-30 minutes post injection during visual stimulation.

Next we attempted to control for the possibility that the differences described between V1 and ACC might be due to differences in the recorded neuronal compartment, or differential effects of MK801 on axons versus somas. We made recordings from the retrosplenial cortex which also provides long distance top-down input to V1, and in which both axons and somas are optically accessible, and found that somas and axons behaved similarly suggesting the effects observed are due to different effects of MK801 on different brain regions (Supplementary Fig. 2).

Previous studies have reported an alteration in inhibitory/excitatory balance in the mPFC following global NMDAR antagonism, resulting from decreased activity of fast spiking inhibitory neurons and increased activity of putative excitatory neurons^5,11^. Given the contrasting pattern of reduced net activity with NMDAR antagonism we observed in V1 we sought to examine what excitatory/inhibitory activity might underlie it. We broke down the population into genetically identified parvalbumin positive inhibitory neurons (PV+; Fig 2a) and putative excitatory pyramidal neurons (Pyr) and found that the activity of the two populations was tightly coupled under control conditions (mean R = 0.81 ± 0.04, n = 7 mice; Fig 2 d,e,f). We then compared the activity of the two populations following MK801 administration and found that the reduction in net population activity was associated with decreased activity of both cell types (Baseline normalised activity: Pyr Saline = 0.96 ± 0.095, PV+ Saline = 0.99 ± 0.109, n = 7 mice; Pyr MK801 = 0.58 ± 0.072, PV+ MK801 = 0.607 ± 0.083, n = 9 mice; Pyr P < 0.01; PV+ P < 0.05; Fig 2f,g). We additionally calculated a normalised ratio of PV+ inhibitory activity to putative excitatory activity and examined signal correlations of the two sub-populations before and after drug treatment, and found that both measures remained stable over the course of the experiment in both treatment groups (Fig 2f,h). These findings together suggest that global NMDAR hypofunction has different consequences on excitatory/inhibitory balance in different cortical regions, resulting in increased activity of top down inputs to sensory cortex but decreased activity of neurons within V1.

**Figure 2:**
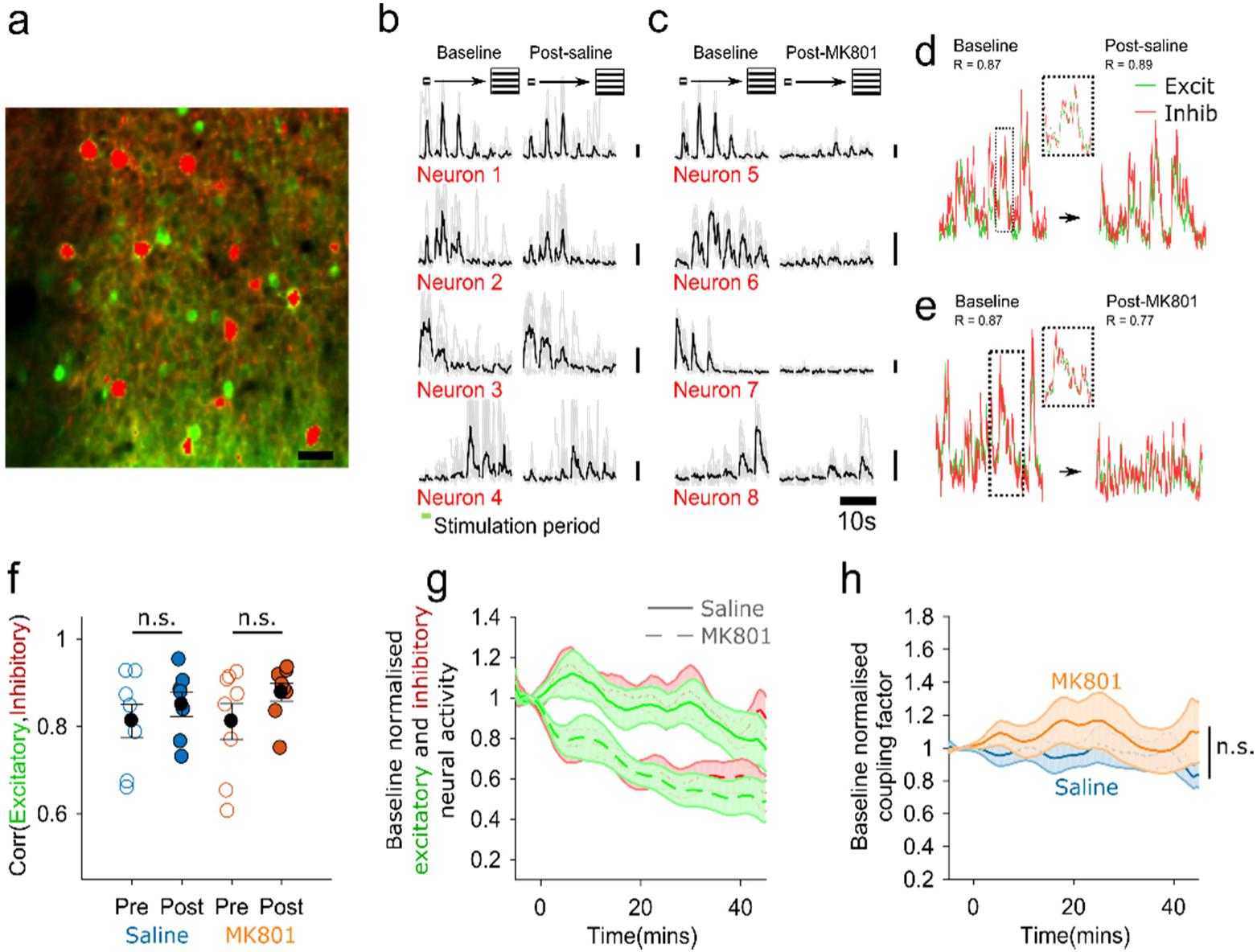
Putative excitatory and inhibitory populations show coupled reduction in activity during NMDAR antagonism. (**a**) PV+ interneurons co-labelled with tdTomato and GCaMP6S. (**b-c**) Example pre and post injection stimulus evoked responses of PV+ interneurons in saline (**b**) and MK801 (**c**) conditions. (**d-e**) Example traces of highly correlated putative excitatory (green) and inhibitory (red) population activity in V1 at baseline and 30 mins after injection of saline (**d**) or MK801 (**e**). (**f**) Correlation of putative excitatory and inhibitory populations in individual animals pre and post injection. (**g**) Timecourse of putative excitatory (green) and inhibitory (red) population activity in saline (solid) or MK801 (dashed) conditions. (**h**) Timecourse of baseline normalised excitatory:inhibitory coupling factor in saline (blue) and MK801 (orange) conditions.

We finally investigated the functional consequences of this altered ratio of top-down verses bottom-up influence on population encoding of sensory stimuli. A multi-class support vector machine (SVM) classifier was used to decode which visual stimulus was presented based on V1 neural activity, and to examine the effect of NMDAR antagonism on decoding accuracy. The SVM was trained on the baseline (pre saline or pre drug) visually evoked activity, and then tested over the following 45 minutes after injection on independent data. We used a leave-n-out strategy to first demonstrate that the SVM could decode V1 activity with a high degree of accuracy during the baseline period, and that decoding accuracy was similar at baseline (pre-treatment) in saline and drug groups (saline: accuracy = 64.1 ± 2.29%, shuffled accuracy = 16.80 ± 0.12%, n = 7 mice; MK801 accuracy = 62.3 ± 4.06%, shuffled accuracy = 16.66 ± 0.11%, n = 9 mice; Fig 3a). We next tested the SVM on a sliding window of 30 visual stimulation trials at a time, normalising classifier performance to the pre-drug/saline period, and observed a decline in classification accuracy after MK801 treatment (accuracy = 41 ± 6.7%, p < 0.001) while classifier performance did not change significantly after saline treatment (accuracy = 56 ± 7.0%, p = 0.28) or in the shuffled condition (Fig 3b,c, see Supplementary Fig. 3 for a similar analysis of decoding of stimulus orientation). In order to assess the effect of differences in mean population activity in the pre and post periods, visual responses were scaled to equalise pre and post administration population response amplitude and a similar decline in classifier performance was observed following MK801 administration (Fig 3c, ‘Post norm’). We next asked whether there is a loss or an alteration of stimulus encoding during NMDAR hypofunction. Instead of training the model on baseline visually evoked activity we divided the post administration period into 4 epochs and performed leave-n-out cross validated training and testing on data from each epoch independently. This resulted in stable classifier performance over the course of the post administration period (Fig 3d) indicating that stimulus encoding is present following MK801 administration but in an altered scheme, while following saline administration the pre-administration scheme is preserved.

**Figure 3:**
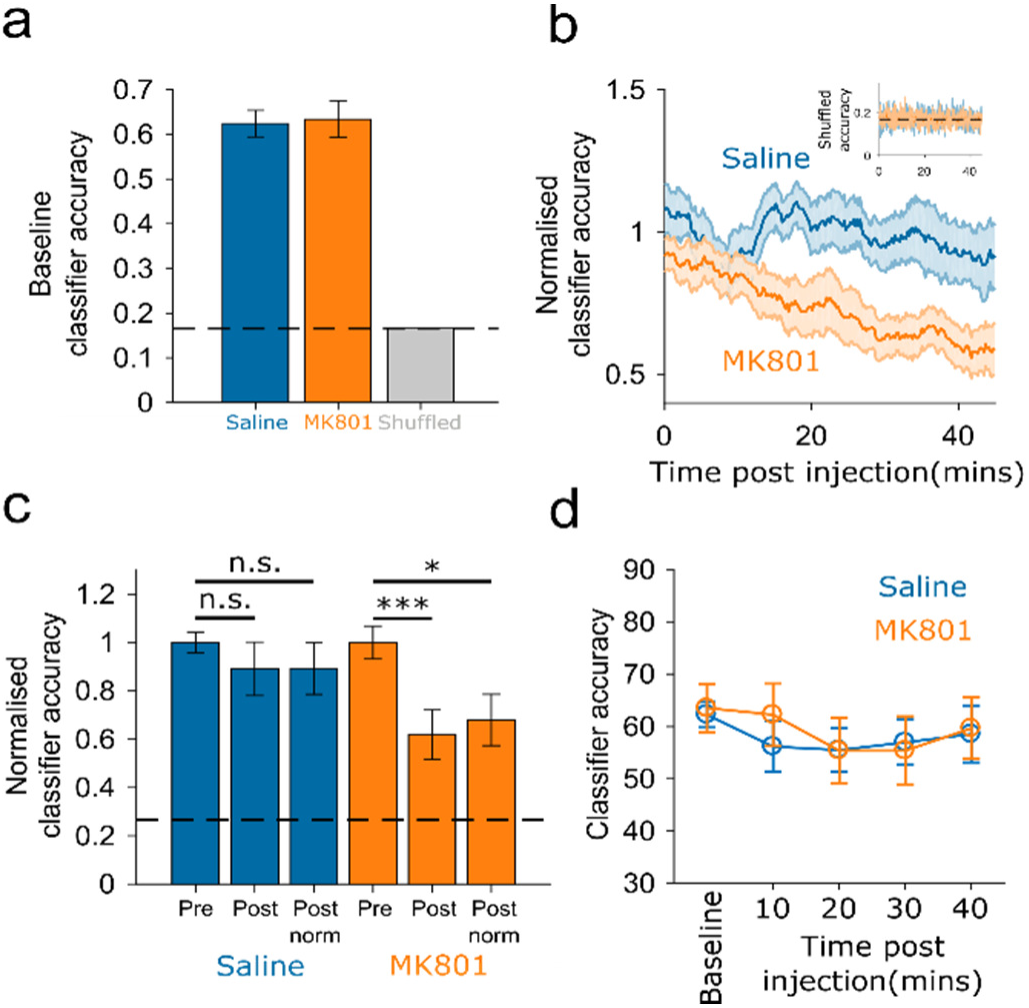
The effects of NMDAR antagonism on support vector machine population decoding of visual stimulus. (**a**) Accuracy of classifier at baseline. (**b**) Timecourse of normalised classification accuracy after training on baseline data, following saline (blue) or MK801 (orange) administration, and timecourse of shuffled performance (inset). (**c**) Average baseline normalised classification accuracy flowing saline (blue) or MK801 (orange) administration, without or with scaling of mean population response amplitude (‘Post’ and ‘Post norm’ respectively). (**d**) Classification accuracy at different time points post injection calculated using ‘leave-n-out’ cross validation when training data is from the same epoch as testing data.

In summary this study provides evidence that global NMDAR antagonism, a pharmacological model of schizophrenia which is known to elicit perceptual disturbances in humans, results in a coordinated reduction in both excitatory and inhibitory activity in V1, in parallel with enhanced top-down drive from the ACC. We suggest that this shift in balance between top-down and bottom-up signals may result in perceptual disturbances by increasing the influence of internally generated signals (representing, for example, prior expectations) over sensory signals from the outside world. This may be one instance of a more general pattern of top-down/bottom-up imbalance in different cortical circuits and at different levels in the hierarchy of cortical processing in neuropsychiatric disease. Given the convergence of genetic risk factors for schizophrenia and related disorders on the NMDA receptor complex and associated synaptic components^4^ these processes may be broadly relevant to the pathogenesis of altered cortical processing in psychiatric disorders.

## Experimental procedures Animals

All experimental procedures were carried out in accordance with institutional animal welfare guidelines, and licensed by the UK Home Office. Experiments were carried out on adult mice (aged >P90). For experiments in which PV interneurons were labelled this was achieved by crossing the B6.Cg-Gt(ROSA)26Sortm14(CAG-tdTomato)Hom/J and B6;129P2-Pvalbtm1(cre)Arbr/J (Jackson Laboratory, JAX Stock#007914 and 008069 respectively). Mice were housed under normal light conditions (14h light, 10h dark) and recordings were made during the light period.

### Animal surgical preparation and virus injection

Aseptic surgical procedures were conducted based largely on previously described protocols^2^. Approximately one hour prior to cranial window surgery and virus injection, animals were administered with the anti-biotic Baytril (5mg/kg, s.c.) and the anti-inflammatory drugs Carprofen (5mg/kg, s.c.) and Dexamethasone (0.15mg/Kg, i.m.). Anaesthesia was induced and maintained using Isoflurane at concentrations of 4%, and 1.5-2% respectively. After animals were stereotaxically secured, the scalp and periosteum were removed from the dorsal surface of the skull, and a custom head plate was attached to the cranium using dental cement (Super Bond C&B), with an aperture approximately centred over the right primary visual cortex, or retrosplenial cortex. For injections into V1 transcranial intrinsic signal imaging was then used to determine the precise location of V1, after which a 3mm circular craniotomy was performed, centred on the area of V1 which responded to visual stimulation at an elevation of 20 deg and azimuth of 30-40 deg. For injections into ACC a small craniotomy was first made over the region (0.3 mm lateral of bregma) either using a dental drill or by thinning the overlying and then piercing a small hole using a hypodermic needle, after which a lager 3mm circular craniotomy was performed over V1. For injections into retrosplenial cortex for retrosplenial soma imaging a 3mm circular craniotomy was performed over the region (centred 0.4 mm lateral and 2.4 mm posterior to bregma). For imaging of retrosplenial cortex axons in V1, a small craniotomy was first made over retrosplenial cortex (centred 0.4 mm lateral and 2.4 mm posterior to bregma) either using a dental drill or by thinning the overlying and then piercing a small hole using a hypodermic needle, after which a 3 mm circular craniotomy was performed over V1. After craniotomy injections of a virus to drive expression of GCaMP6S (AAV1.Syn.GCaMP6s.WPRE.SV40; titre after dilution 2x10^11^ GC/ml) were made into the relevant region (V1, depth = 250µm, 40nl at 1-3 sites; ACC, depth = 800µm, 100nl at 1 site; RCx for soma imaging, depth = 250µm, 40nl at 1-3 sites; RCx for axon imaging, 40nl at 1 site). Injections were made using a microsyringe driver (WPI, UltraMicroPump) coupled to a pulled and bevelled oil filled glass micropipette with a tip outer diameter of approximately 30μm. After injection the craniotomy was closed with a glass insert constructed from 3 layers of circular no 1 thickness glass (1x5 mm, 2x3 mm diameter) bonded together with optical adhestive (Norland Products; catalogue no. 7106). After surgery animals were allowed at least 2 weeks to recover after which they were either habituated to head fixation passively or during a visual discrimination task.

### Imaging and locomotor behaviour

*In vivo* 2-photon imaging was performed using a resonant scanning microscope (Thorlabs, B-Scope) with a 16x 0.8NA objective (Nikon). GCaMP6 and tdTomato were excited at 980nm using a Ti:sapphire laser (Coherent, Chameleon) with a maximum laser power at sample of 50mW. Data was acquired at approximately 60Hz and averaged, resulting in a framerate of approximately 10Hz. Cortical surface vascular landmarks were used to locate the same neurons between sessions. During 2-photon imaging animals were free to run on a custom designed fixed axis cylindrical treadmill, and movement was measured using a rotary encoder (Kübler, 05.2400.1122.0100). Imaging, behavioral and visual stimulation timing data were acquired using custom written DAQ code (Matlab) and a DAQ card (NI PCIe-6323, National Instruments).

*In vivo* intrinsic signal imaging was performed using previously described methods^3^ using either a custom built system based around a MAKO G-125B camera (AVT) or a commercially available system (Imager 3001, Optical Imaging Inc.)

### Visual stimuli

For V1 recordings the preferred population retinotopic location of the field of view of neurons was determined in advance using circular 30x30 deg drifting horizontal gratings with temporal frequency of 2 Hz and spatial frequency of 0.05 cycles per degree. Each stimulus appeared and was stationary for 5 seconds, drifted for 2 seconds, was stationary for 2 further seconds and then disappeared. Trials were spaced by 3 seconds, during which a grey screen was displayed. Visual stimuli were generated using the psychophysics toolbox^4^, and displayed on calibrated LCD screens (Iiyama, BT481). Having established retinotopic preference, orientation tuning was next measured using circular gratings with the same temporal and spatial frequency, at the identified preferred location, and displayed at 12 different orientations. For recordings of ACC axons in V1, RCx somas or RCx axons in V1 the visual stimulus was positioned in the binocular area directly in front of the animal.

### Experimental design

In order the measure the effect of NMDAR blockade a 6 minute recording of baseline activity was made (either in V1 somas, ACC axons in V1, RCx somas or RCx axons in V1) during which the animal was exposed to horizontally oriented grating stimuli with the spatial and temporal frequency and position determined as described above, which varied in size between 10-60 degrees in steps of 10 degrees. After this baseline period the recording and visual stimulation was briefly paused, animals were injected with either saline or MK801 (dose), after which the recording and visual stimulation was resumed for 45 minutes.

### Calcium imaging data analysis

Brain motion was first corrected for using an automated registration algorithm^5^ implemented in Matlab, and data from the pre-injection administration period for registered to the post-injection period. A 20μm border was removed from all frames (more than the maximum brain movement observed) to ensure that all pixels were present in all frames in both sessions. Soma regions of interest were identified using a custom written semi-automated algorithm based on grouping of pixels with correlated time-courses. Pixels within each region of interest were then averaged and background fluorescence contamination was estimated from a 30μm circular area surrounding each soma ROI (excluding other ROIs) and subtracted from the soma ROI signal with a weighting of 0.7. Only cells with somas which were >5% brighter than surrounding neuropil were included in further analysis. For analysis of axonal data, labelled axons were first identified by thresholding the averaged movie frame for the ROI (90% of the 90^th^ percentile value), and then averaging the time courses of the detected pixels. The time series of each ROI was then converted from a raw fluorescence value to dF/F with the denominator F value calculated as the 5^th^ percentile of the smoothed raw trace. Population activity was calculated as the mean timecourse of all detected neurons in a field of view smoothed with a 2.5 minute sliding window. Correlation of excitatory and inhibitory populations were calculated by averaging the time courses of all identified excitatory and inhibitory neurons and then calculating the Pearson correlation coefficient in the baseline and post-saline/drug period. The timecourse of the coupling of excitatory and inhibitory populations was calculated by first smoothing the traces of both cell types with a sliding averaging window (window size of 5 secs), then normalising each trace to the baseline period (to control for baseline differences in levels of activity in the excitatory/inhibitory populations), and then dividing the excitatory trace by the inhibitory trace at each time point. In order to identify tdTomato labelled PV+ neurons, cells were semi-automatically classified based on a thresholded mean registered red channel image, which was eroded and then dilated to remove small areas of labelling of neural processes, after which classification was verified manually.

### Support Vector Machine

The Support Vector Machine analysis was implemented in Matlab using the LIBSVM library with radial basis function kernel type and C-SVM multi-class classification options^6^. Visual stimulus identity was decoded from the responses of the entire population of recorded neurons from each animal, where individual neurons were considered as features. In order to calculate population decoding accuracy at baseline (Fig 3a), or post injection epoch-by-epoch analysis (Fig 3d), a “leave-n-out” strategy was used to calculate and test the multiclass SVM, whereby the model was repeatedly trained on all but 10% of randomly chosen trials and then tested on the remaining trials. In order to test classification accuracy in the post-drug/saline period the classifier was trained on all trials from the baseline period, and then tested on a sliding window of 30 independent trials, advancing in 1 trial steps, from the post saline/drug administration period. This process was performed either on unmodified response amplitude data or on response amplitudes which had been normalised such that mean population response amplitudes in the pre and post injection conditions were equal. Performance was then normalised to the cross validated estimate of baseline classification accuracy. The SVM classification accuracy timecourse traces were then averaged across animals. Chance classification accuracy was calculated by testing the SVM model with shuffled stimulus identifiers.

## Acknowledgements

### Author contributions

A.R. and J.H. designed experiments. A.R. performed imaging experiments and analysed data. A.R., E.B. and A.P. performed surgeries. A.P. and E.B. provided histology data. E.B. and A.R. developed the anterior cingulate cortex imaging protocol. A.P. and A.R. developed the retrosplenial cortex imaging protocol. A.R. and J.H. wrote the paper.

## Acknowledgments

This work was supported by a Wellcome Trust Strategic Award (503147) to Michael J Owen, JH, Lawrence Wilkinson, Adrian Harwood, Meng Li, David Linden, John Aggleton, Vincenzo Crunelli and Derek Jones and a Wellcome Trust ISSF Seedcorn Award to AR (508353). For the use of GCaMP6S we acknowledge Vivek Jayaraman, Rex A. Kerr, Douglas S. Kim, Loren L. Looger, Karel Svoboda from the GENIE Project, Janelia Farm Research Campus, Howard Hughes Medical Institute. We thank Chris Burgess for building parts of the data acquisition system, and Kenneth Harris for developing parts of the ROI detection pipeline. We thank Stuart Greenhill and Frank Sengpiel for helpful comments and discussion of the manuscript.

### Competing financial interests

The authors have no competing financial interests.

## Supplementary Materials

**Supplementary Figure 1:**
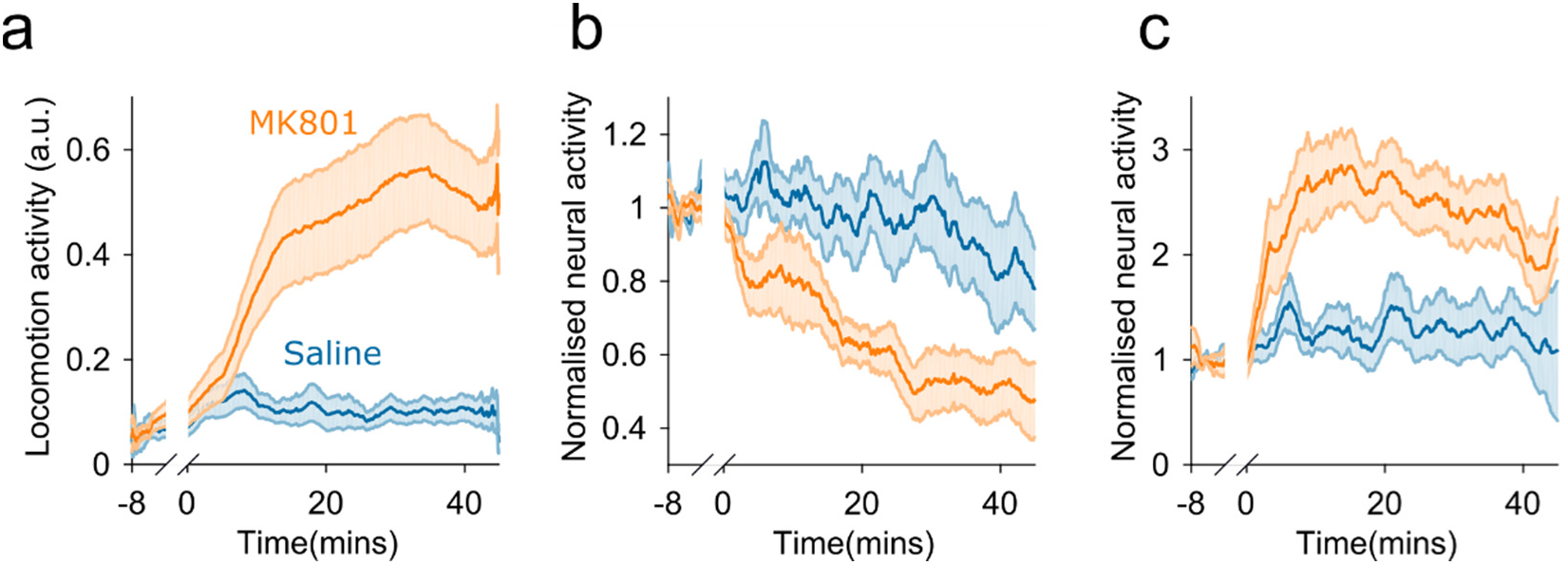
Hyper locomotion behaviour. (**a**) Hyper locomotion observed following MK801 administration. (**b-c**) As in Figure 1e and Figure 1f, but with analysis of V1 (**b**) and ACC axon (**c**) activity limited to periods of locomotion.

**Supplementary Figure 2:**
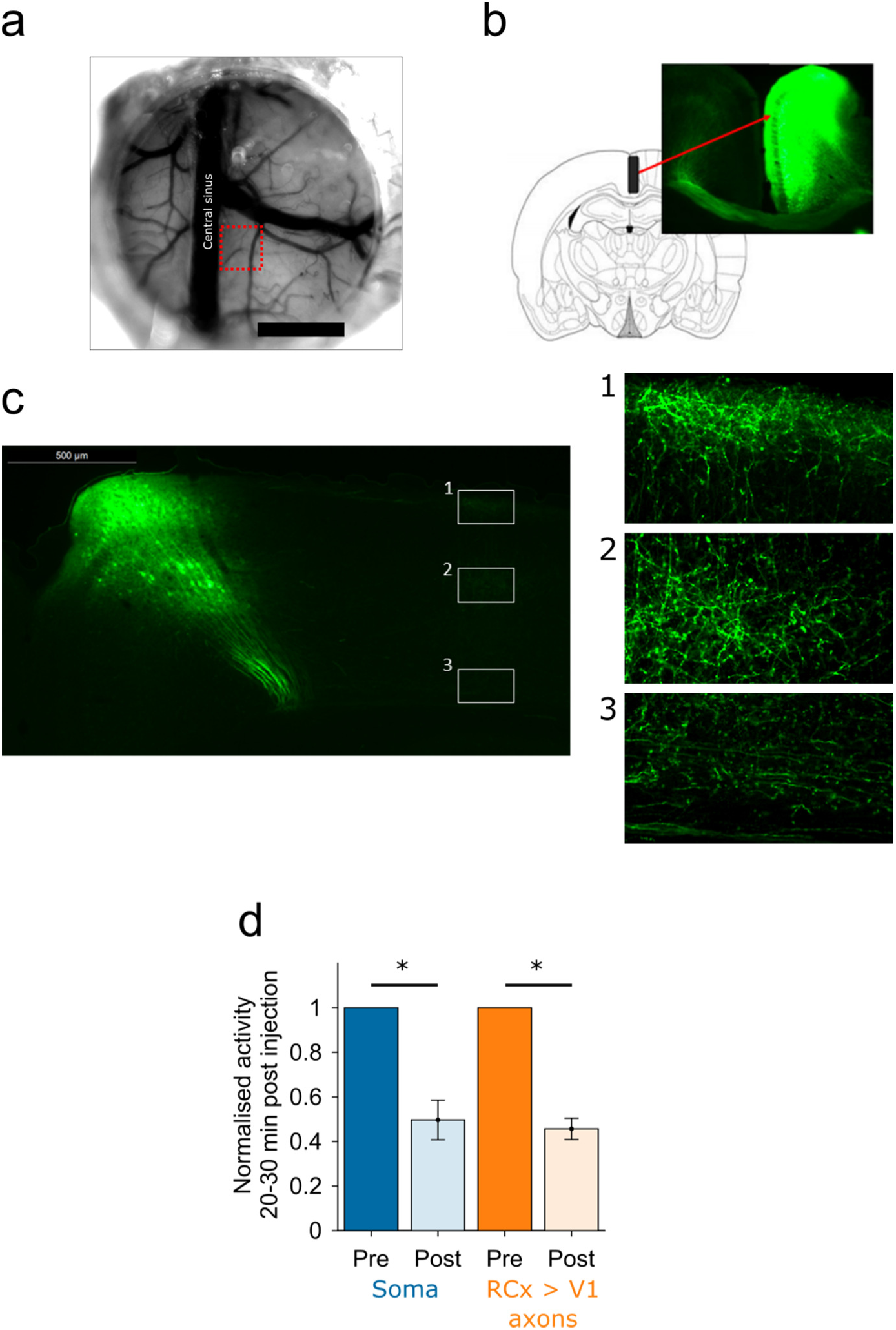
Effects of NMDAR antagonism on somas and axons of neurons from the same population in retrosplenial cortex (RCx). We sought to control for the possibility that differences between the activity of axons from ACC and somas in V1 might be due to the neuronal compartment recorded from (somas vs. axons) rather than brain area differences. We examined this question by recording from the retrosplenial cortex which provides top-down input to the visual cortex and has somas which are optically accessible for 2-photon microscopy. This allowed quantification of another source of axonal top-down input to V1, as well as direct measurement of somas in the region from which the axons originate. This showed firstly that contrary to previous reports *in vitro*^1^, *in vivo* retrosplenial cortical activity is also strongly attenuated by MK801, and secondly that this is reflected in the calcium signals observed in the axons originating in the region. (**a**) Cranial window positioned over RCx. (**b**) Injection site and GCaMP6S expression 3 weeks post injection. (**c**) RCx innervation of visual cortex. (**d**) Normalised activity of RCx somas (blue) and RCx→V1 projecting axons (orange) following MK801 administration, showing similar pattern of reduced activity (baseline normalised activity: Somas = 0.49 ± 0.09, P = 0.03, n = 3 mice, Axons = 0.46 ± 0.05, P = 0.007, n = 3).

**Supplementary Figure 3:**
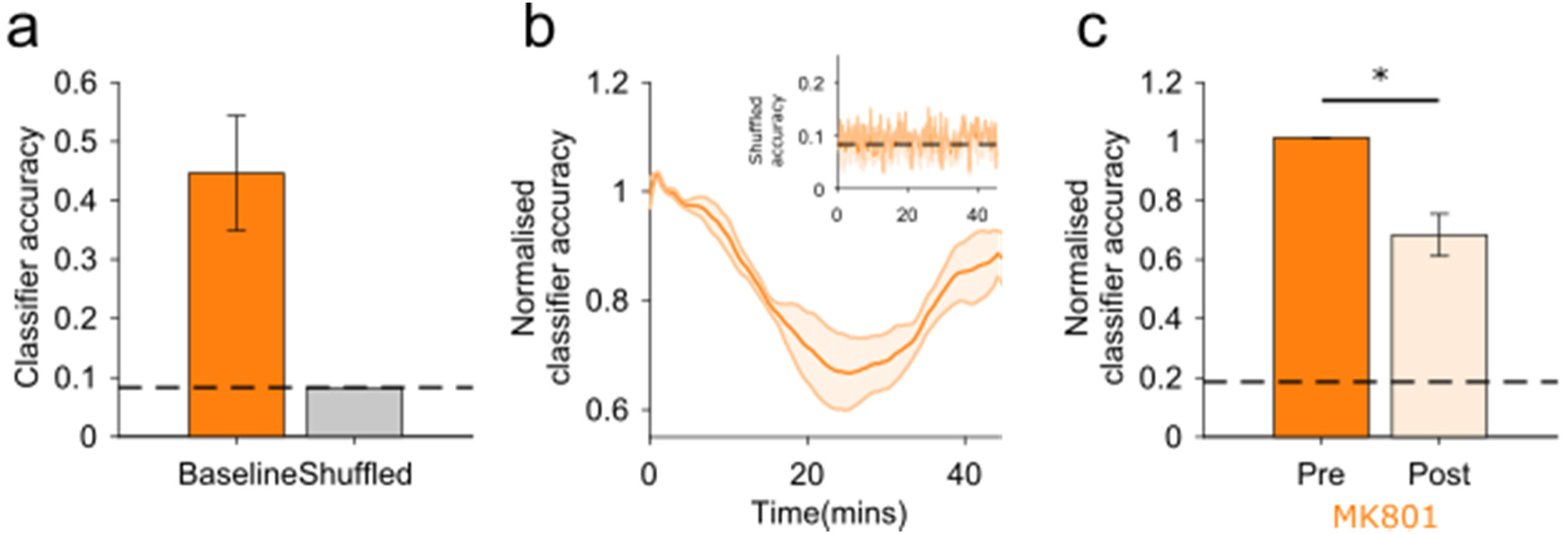
Support vector machine decoding of stimulus orientation. (**a**) Baseline period support vector machine decoding accuracy compared to chance and shuffled (MK801 accuracy = 45.5 ± 0.1%, shuffled accuracy = 8.3 ± 0.1%, n = 3 animals). (**b**) Timecourse of classification accuracy following MK801 administration, and timecourse of shuffled performance (inset). (**c**) Average baseline normalised classification accuracy at 20-30mins post MK801 administration (normalised classification accuracy 68.2 ± 0.1 %, p = 0.03, n = 3 animals).

**Supplementary Video 1: Timelapse video of activity of somas in V1 and ACC→V1 axons over the 45 minutes following MK801 administration.**

